# Pupil-based states of brain integration across cognitive states

**DOI:** 10.1101/2020.12.15.422870

**Authors:** Verónica Mäki-Marttunen

## Abstract

Arousal is a potent mechanism that provides the brain with functional flexibility and adaptability to external conditions. Within the wake state, arousal levels driven by activity in the neuromodulatory systems are related to specific signatures of neural activation and brain synchrony. However, direct evidence is still lacking on the varying effects of arousal on macroscopic brain characteristics and across a variety of cognitive states in humans. Using a concurrent fMRI-pupillometry approach, we used pupil size as a proxy for arousal and obtained patterns of brain integration associated with increasing arousal levels. We carried out this analysis on resting-state data and data from two attentional tasks implicating different cognitive processes. We found that an increasing level of arousal was related to a non-linear pattern of brain integration, with increasing brain integration from intermediate to larger arousal levels. This effect was prominent in the salience network in all tasks, while other regions showed task-specificity. Furthermore, task performance was also related to arousal level, with accuracy being highest at intermediate levels of arousal across tasks. Taken together, our study provides evidence in humans for pupil size as an index of brain network state, and supports the role of arousal as a switch that drives brain coordination in specific brain regions according to the cognitive state.

## 1. Introduction

Arousal-related neuromodulatory systems are a major drive that modulate the brain and give great functional flexibility beyond a relatively fixed structural core. Understanding the mechanism by which this occurs is crucial for understanding how the brain rapidly adapts to different situations, and how a dysbalance in arousal levels may lead to aberrant mental states or conditions (Atzori et al., 2016), such as autism (Anderson et al., 2013), psychosis (Mäki-Marttunen et al., 2020a) or post-traumatic stress syndrome (Pitman et al., 2012).

Sleep, awake or stressed states constitute largely distinguishable arousal states. Within those states, however, it is plausible to think of a continuum of arousal. The fluctuations of arousal within active wake states have been assessed by measuring baseline pupil size, a surrogate measure of arousal. Such studies have shown that baseline pupil size allows distinguishing active task solving versus mind wandering (Kristjansson et al., 2009; Smallwood et al., 2011; Unsworth & Robison, 2016, 2018; van den Brink et al., 2016a), exploitation versus exploration states (Jepma & Nieuwenhuis, 2011; Murphy et al., 2011; Pajkossy et al., 2017), focused attention versus fatigue (Nishiyama et al., 2007) or stress (Maran et al., 2017), all of which may alternate even during the course of a task. In addition to studies in humans, animal studies have shown that states associated with different level of arousal during the wake state show different pupil signatures, as well as different signatures of activity at the neuronal level. For instance, locomotion or whisking is associated to larger pupil sizes as compared to standing still (McGinley et al., 2015a; McGinley et al., 2015b; Reimer et al., 2014; Stitt et al., 2018). Thus, it is possible to describe arousal variation by looking at pupil size behavior while comparing tasks associated to different cognitive processes or varying engagement to external demands.

To face present demands, the brain has the ability to adjust the degree of interaction between its constituting neural systems (Shine & Poldrack, 2018). This ability is ensured in part by the arousal-related neuromodulatory systems (Guedj et al., 2017a; Shine, 2019; Zagha & McCormick, 2014). These systems, such as the noradrenergic, dopaminergic or cholinergic systems, rapidly modify synaptic properties at more local or more global levels and facilitate the passing of information in neural circuits transmitting relevant information (Arnsten et al., 2012; Krichmar, 2008; Thiele & Bellgrove, 2018). At a macroscopic scale, this may be seen as a reorganization of brain networks interaction, where the flexible balance between integration and segregation of different regions gives rise to a diversity of cognitive states (Shine et al., 2018a). Work using functional magnetic resonance imaging (fMRI) has studied how the brain oscillates between different modes characterized by different degrees of integration between constituting networks (Fukushima et al., 2018). In general, more integrated states have been related to more cognitively demanding tasks (Shine, 2019). However, the expression of brain integration states along a range of arousal and cognitive states have remained largely unexplored.

Previous studies attempted to assess this question by pharmacologically manipulating the arousal-related neuromodulatory systems and measuring brain connectivity. They showed that boosting the catecholamine system (i.e. noradrenaline and dopamine) leads to a rearrangement of the patterns of interaction between brain networks, and that this depends on behavioral state (Eldar et al., 2013; Guedj et al., 2017b; Hernaus et al., 2017; Shine et al., 2018b; van den Brink et al., 2016b). In an fMRI-pupillometry study, Shine et al. found that the temporal course of pupil size covaries with a measure of brain integration during resting state (Shine et al., 2016). Here we argue that to obtain a more complete picture of how arousal adaptively modulates brain interaction, it is necessary to examine a variety of cognitive tasks, capture a range of task engagement or arousal level, and measure the degree of interaction between brain networks.

In this study, we employed pupil size distribution as a surrogate of arousal, and investigated the associated patterns of integration between brain regions. The advantage of using pupil size as an index of arousal is that it allows analyzing a physiological range of arousal levels, that is, without external (e.g. pharmacological) manipulation. We further analyzed tasks involving different cognitive processes or engagement to external demands. The goal of the present study was to assess the topological brain signatures of increasing arousal level, and whether they differed across tasks. The tasks compared were: resting state, a continuous performance task (which requires sustained monitoring), and a multiple-object tracking task. We expected that varying levels of effort or engagement in the tasks would consequently span a range of arousal levels (Eckstein et al., 2017). We hypothesized that larger arousal during tasks would be related to more brain integration (Shine et al., 2016), and that the brain patterns at increasing arousal would be specific of cognitive states, that is, resting state vs. attentional tasks. We also inspected the relation between behavior and arousal in our two tasks.

Our results showed that an increasing level of arousal was related to an inverted-U pattern of brain integration in specific regions. The salience network consistently showed this pattern across tasks, indicating a generalized effect of arousal on this network. Other areas varied their level of integration in specific cognitive states. We also found higher accuracy at intermediate levels of arousal in the two tasks. This provides new evidence in humans that the dynamic recruitment of arousal-related systems has a precise brain signature, which is to coordinate the communication across brain networks in a task-specific manner. This “switch” of the brain may be engaged flexibly in the context of different external demands.

## 2. Experimental procedures

### 2.1 Dataset

60 participants (age: 27 ± 5, range: 20-42, N female = 39) were recruited in the University of Oslo. The work was carried out in accordance with the Declaration of Helsinki. Informed written consent was obtained in accordance with the protocols approved by the local Ethics Committee, and all participants were given 100 NOK per hour for their participation. None of the participants had a present or past history of serious disorders. Five participants were excluded from the functional analysis because the pupil data was absent.

### 2.2 Tasks

The participants were asked to perform three different tasks in the scanner. A resting state (RS) scan was acquired, where participants had to lie still and look at a fixation cross in the middle of the screen. This task lasted 10 minutes.

In addition, participants performed two runs of the AX-continuous performance task (AX-CPT). This task is commonly used to study cognitive control (Mäki-Marttunen et al., 2019; Servan-Schreiber et al., 1996). Each trial of the standard AX-CPT consisted of two displays: first, a contextual cue was presented, and after a delay, the probe followed. Participants were instructed to detect target trials in which the cue was a prespecified stimulus (“A”), and the probe was another prespecified stimulus (“X”). Participants had to identify target trials by pressing the appropriate button (i.e. target response). For all other pairs (non-“A”, and/or non-“X”), a non-target response should be given. The “AX” target pair constituted the majority of the trials (64% of the trials), while the other three trial types were relatively infrequent (12% each) (Servan-Schreiber et al., 1996). For stimuli, we used human silhouettes. In each trial, cues were presented for 300 ms followed by a jittered delay period (mean of 3000 ms; range: 2500 - 3500 ms). Then the probe was presented for 1200 ms. Participants could respond during this period, which was followed by a feedback message (“Correct” or “Wrong”, 500 ms). Trials were separated by a 2000 ms interval with a fixation cross (range 1500 - 2500 ms). Participants performed 50 trials per run (100 trials in total). Within each run, trials were blocked in two blocks of 25 trials, and each block was followed by a 18 sec resting block. For further details on the design and display, please refer to our previous study (the current design corresponds to the low load condition of Mäki-Marttunen et al. 2019).

Lastly, participants performed three runs of the multiple-object tracking (MOT) task. In each trial of the MOT task, participants were presented with 10 blue rounded objects. A number of these objects were then cued as targets by turning red for 2.5 secs. Afterwards, these targets turned back to blue, and then all the disks started to move independently in random directions. The participant’s task was to track the objects indicated as targets as all the objects moved around. The tracking period lasted 12 secs. At the end of the trial, one object was highlighted and participants had to indicate whether this was a target or not. The load of the task was varied by increasing the number of objects to track: two, three, four or five objects. In addition, some trials were passive, were participants observed the ten objects moving, but none of them was indicated as target. Each run consisted of 25 trials in total (five of each load level and five of passive viewing). The task was adapted form Alnæs et al. 2014.

### 2.3 Pupillometry

For the acquisition of pupil data we used an MRI-compatible EyeLink 1000 eye tracker. The sampling rate was 1000Hz. The pupil data followed standard preprocessing, with missing data and blinks interpolation, band-pass filtering (0.01-10Hz) (code adapted from Urai et al. 2017) and resampling to the number of MRI volumes. For the pupil analysis, not all concurrent pupil data could be acquired or was usable for each task. The final sample size for each task was: resting-state: N = 49; AXCPT: N = 53; MOT: N = 49. The pupil size distribution from a sample subject is displayed in **Figure 1.b**. The demeaned pupil size vectors were used to compute a distribution of pupil values (**Figure 1.c**). Based on this distribution, five intervals or bins of pupil size were established: [−400, −200], [−200, 0], [0,200], [200−400], and [>400] (see Stitt et al. 2018 for a similar approach). Any value under −400 was set to zero because this range was likely to contain blink or edge (e.g. initial dip) artifacts.

**Figure 1.**
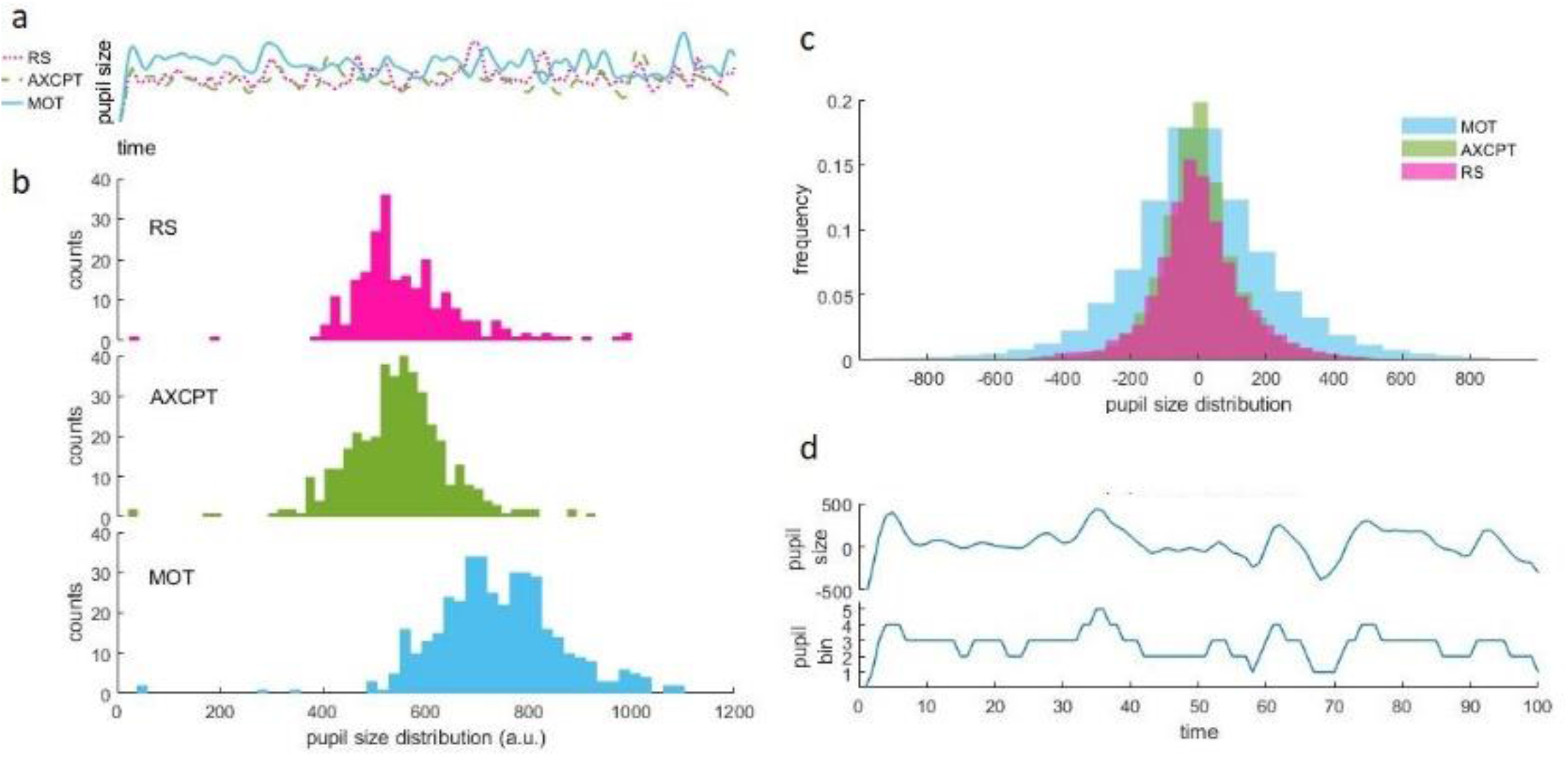
**a:** Sample timecourses of raw pupil size values in one participant, and **b**: the corresponding distribution, separated by task. **c**: Group distribution of demeaned pupil size values across all tasks. **d**: example of a pupil size time course in one run and its corresponding vector of assigned pupil bin (1 corresponds to intervals with smallest pupil, 5 corresponds to largest pupil).

Based on this categorization, the time points of individuals’ pupil vectors were assigned a pupil bin (see **Figure 1.d** for an example). In the supplementary material, we provide further characterization of the pupil size bins.

### 2.4 Image acquisition and preprocessing

MRI sessions took place in the 3T Philips Achieva scanner (8-channel head coil) at Rikshospitalet, Oslo. Each scanning session started with an anatomical scan. A 10-minute resting state protocol consisting of 250 volumes was collected. Whole-brain functional images were acquired using a spin-echo echo-planar (EPI) sequence sensitive to blood-oxygen-level-dependent (BOLD) magnetic susceptibility (TR = 2500 ms; Flip angle = 90°; number of slices: 42; voxel size: 3 mm^3^). During the resting state, a cross was projected on a screen positioned at the head end of the litter. Participants viewed the screen through a mirror placed on the head coil.

For the task scans, EPI sequences were acquired (TR = 2208 ms; Flip angle = 90°; number of slices: 42; voxel size: 3 mm^3^). Each AXCPT run consisted of 212 volumes, and each MOT run, of 225 volumes. Responses were given through buttons on an fMRI-compatible joystick device, with one button corresponding to target response (AXCPT) or target ball (MOT), and another to non-target response.

The functional images of each subject were first visually inspected for anomalies and then submitted to a standard preprocessing pipeline using SPM 12 implemented on MATLAB (Mathworks). Images were first corrected for time delays and realigned using 6 parameters of movement. The data were normalized to a standard template in the MNI system (image size: 75×95×75, voxel size: 2×2×2 mm3) using the unified segmentation algorithm of SPM 12 with six tissue maps as priors. Images were then smoothed using 8 mm FWHM Gaussian kernel. Structural and functional images were loaded in the CONN toolbox (Whitfield-Gabrieli & Nieto-Castanon, 2012). Further preprocessing steps included band-pass filtering (0.008-0.1 Hz), linear detrending and denoising. For denoising, the five main principal components of the signal from white matter and cerebro-spinal fluid were included as confounds. For the first level design, the imaging data was modeled with the pupil interval vectors of each run. For each subject, functional connectivity was calculated as the Pearson correlation between 30 regions of interest (ROIs) that are part of seven canonical resting state networks (Default Mode, Sensory-Motor, Visual, Salience/Opercular, Dorsal Attention, Fronto-parietal and Language). The ROIs are available within the CONN toolbox (www.nitrc.org/projects/conn, RRID:SCR_009550). The average connectivity matrix for each subject and each pupil category was obtained. Based on the correlation matrices, a measure of brain integration, participation coefficient (PC, Bertolero et al. 2017), was calculated. This measure is based on graph theory and describes the degree in which brain regions, or nodes, interact with other regions outside its module or network (Medaglia, 2017; Rubinov & Sporns, 2010):

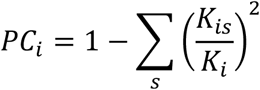

where *K_i_* is the sum of node *i*’s edges and *K_is_* is the sum of *i*’s edges to community *s*. The PC has been previously used to study the state of integration of the brain, in particular in relation to arousal fluctuations (Shine et al., 2018a; Shine et al., 2016; Shine et al., 2018b). In addition, we calculated the segregation index (SI), which is calculated based on the relation between within and between network connectivity for each brain network:

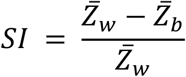

where 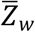 is the z-transformed mean correlation between nodes of a given network and 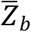 is the z-transformed mean correlation of the nodes of the network with nodes outside the network. The segregation index, as opposed to the participation coefficient, is a measure of how segregated are brain systems (Chan et al., 2014).

### 2.5 Statistics

To compare the participation coefficient values for the different networks and pupil bins, we used a linear mixed model. The reason is that not all pupil bins were present in some participants, and this type of model allows to deal with missing data for some levels in a factor. Statistical analyses were run in SPSS (IBM). Pupil bins (levels 1 to 5) were designated as the fixed factor, and the PC constituted the dependent variables. For correction of multiple comparisons, an FDR correction (Benjamin-Hochberg threshold of 0.2) was applied. In addition, linear mixed models were run for behavioral data of each task (accuracy and reaction times) and SI, with pupil bin as the fixed factor.

## 3. Results

This study pursued the question of whether increasing arousal was related to specific patterns of brain interaction, with either task-independent (that is, common across tasks) and task-dependent aspects. For this, we defined intervals based on the pupil size distribution and examined the brain patterns of integration in each interval and each task.

As a measure of brain integration, we calculated the PC of thirty regions of interest (ROIs) grouped in seven canonical networks: Default Mode (DMN), Sensory-Motor (SMN), Visual (VIS), Salience/Opercular (SAL), Dorsal Attention (DAN), Fronto-parietal (FPN) and Language (LAN). A set of regions in each task showed a significant relation between their degree of integration and pupil bin (**Figure 2, Table 1, Figure S2**). Across tasks, integration in the salience network consistently appeared as significantly associated with bin of pupil size. Other areas showed a significant trend only in one of the tasks.

**Figure 2.**
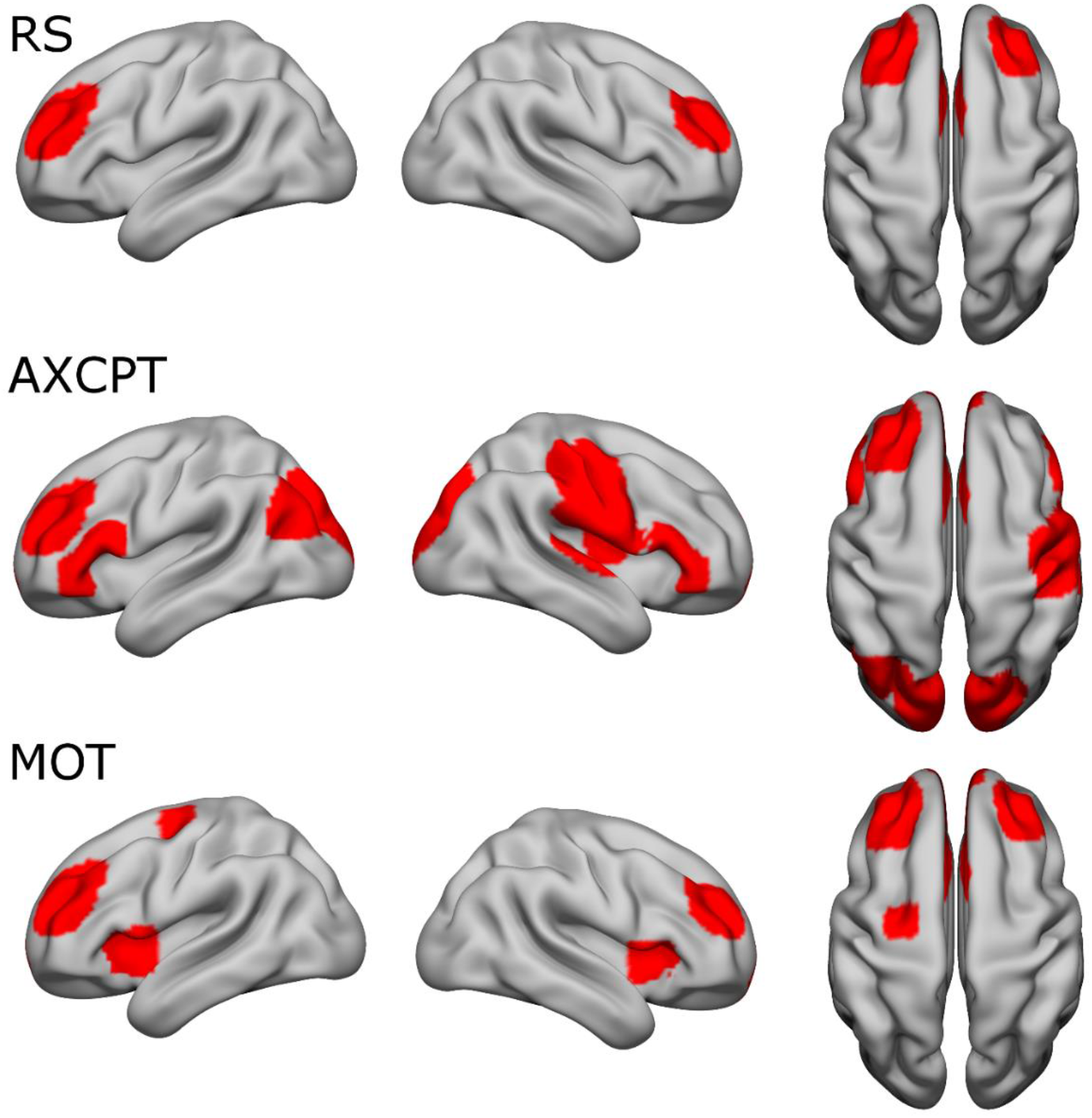
Brain regions for which the participation coefficient showed a significant relation with pupil size bin.

**Table 1.** Brain regions and corresponding network for which the participation coefficient showed a significant relation with pupil size bin. Only regions surviving multiple comparisons (FDR) correction within each task are shown.

| TASK | NETWORK | REGION | P VALUE |
| --- | --- | --- | --- |
| RS | SAL | ACC | 0,013 |
| RS | SAL | r RPFC | 0,013 |
| RS | SAL | l RPFC | 0,014 |
| AXCPT | DMN | MPFC | 0,001 |
| AXCPT | DMN | l LP | 0,001 |
| AXCPT | SAL | ACC | 0,015 |
| AXCPT | SAL | l RPFC | 0,017 |
| AXCPT | SMN | r Lateral | 0,016 |
| AXCPT | LAN | l IFG | 0,019 |
| AXCPT | LAN | r IFG | 0,02 |
| MOT | SAL | l RPFC | 0,001 |
| MOT | SAL | ACC | 0,002 |
| MOT | SAL | l Alnsula | 0,003 |
| MOT | SAL | r Alnsula | 0,015 |
| MOT | SAL | r RPFC | 0,029 |
| MOT | DAN | l FEF | 0,003 |
| MOT | DMN | MPFC | 0,015 |

To see whether pupil size was associated to an increase or decrease of regions’ integration, we inspected the relation between PC and pupil size in each task. We observed an inverted-U pattern, with larger brain integration at low and high pupil size bins in all three tasks (**Figure 3**).

**Figure 3.**
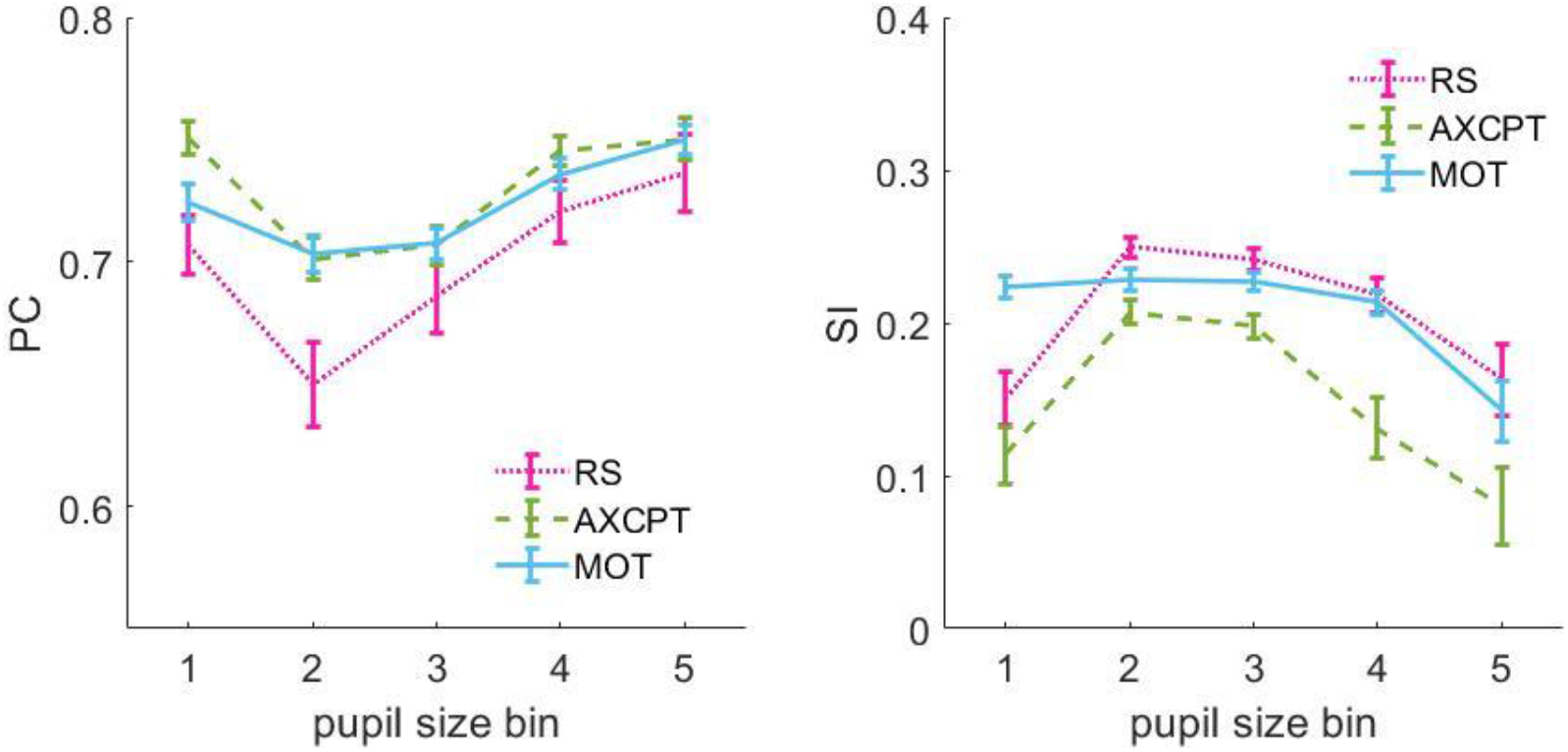
Average trend of the brain regions that showed a significant relation between participation coefficient (our measure of brain integration) and pupil size bin (**Table 1**) per task. Pupil size bin 5 corresponds to largest pupil values.

We complemented the analysis of brain integration with a network-specific index of segregation (SI). This index showed a significant effect of pupil bin in several networks in each task (**Table 2, Figure S3**). The SI of the salience network showed a significant effect of pupil bin across tasks, while other networks showed a significant effect in some of the tasks. The relation between SI and pupil bin in RS and AXCPT followed a U pattern, with more segregation at intermediate pupil size bins (**Figure 3**), while for MOT there was decreased SI only at the largest pupil bins.

**Table 2.** Segregation index per network and task. *: significant effect of pupil size bin after Bonferroni correction.

| SI | RS | AXCPT | MOT |
| --- | --- | --- | --- |
| DMN | 0,001* | 0,001* | 0,157 |
| SMN | 0,001* | 0,001* | 0,014 |
| VIS | <0.001* | 0,026 | 0,493 |
| SAL | 0,007* | <0.001* | <0.001* |
| DAN | 0,002* | 0,02 | <0.001* |
| FPN | 0,002* | <0.001* | 0,009 |
| LAN | 0,022 | 0,188 | 0,911 |

Finally, we inspected whether behavior was related to pupil size bin. We observed a significant effect in accuracy in both AXCPT and MOT tasks (AXCPT: F = 7.567, p <0.001, MOT: F = 9.022, p <0.001), and in MOT, we also observed a significant effect in the reaction times (AXCPT: F = 2.356, p = 0.056, MOT: F = 9.201, p <0.001). The variables followed a U-shaped curve, reaching a maximum at intermediate levels of pupil size (**Figure 4**).

**Figure 4.**
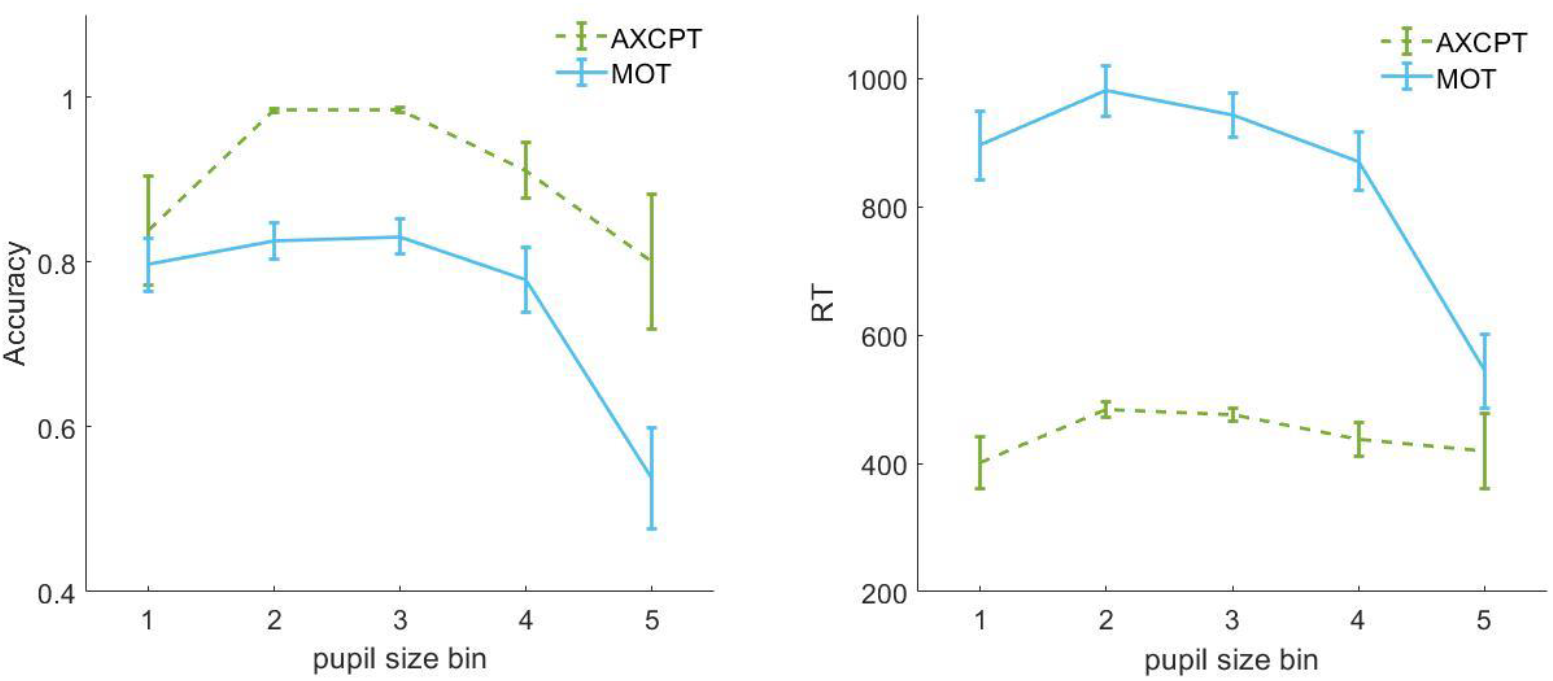
Average accuracy and reaction time per task as a function of pupil size bin.

## 4. Discussion

In the present work, we defined levels of arousal based on pupil size and examined the patterns of brain integration across arousal levels for three different tasks: resting state, a sustained attention task (AXCPT) and a visual attention task (MOT). We found that specific areas showed a U-pattern, with low integration at intermediate levels of arousal and an increase towards larger arousal in the different tasks. The analysis with a measure of segregation showed a complementary pattern, with higher segregation at intermediate levels. Our results support the view that arousal-related neuromodulatory systems exert a generalized effect in brain coordination. Moreover, we provide evidence in humans of the brain network patterns that shift their degree of integration with arousal across mental states.

The first step in this study consisted on identifying pupil size-based levels of arousal. Ocular measures have been used as a continuous measure of arousal in animals (Chang et al., 2016; Reimer et al., 2014; Wang et al., 2016), and pupil size has been widely employed as a measure of arousal and task engagement in cognitive experiments (Eckstein et al., 2017; van der Wel & van Steenbergen, 2018). In support of this, neuronal firing in one of the main neuromodulatory centers that regulate effort and arousal, the locus coeruleus (LC, Aston-Jones and Cohen 2005; Berridge and Waterhouse 2003; Carter et al. 2010; Costa and Rudebeck 2016; Samuels and Szabadi 2008), located in the brainstem, strongly predicts pupil dilation (Joshi et al., 2016; Rajkowski, 1993). Indirect evidence of the link between LC activity and pupil dilation in humans was provided by fMRI studies (Murphy et al., 2014; Mäki-Marttunen & Espeseth, 2020). More recently, studies in rodents have shown that different distributions of pupil size values characterize different behavioral states, with more active states (e.g. walking or whisking) peaking at larger pupil values than the passive states, (e.g. stillness, McGinley et al. 2015b; Reimer et al. 2014). In other words, behavioral states associated with different levels of arousal and LC activity are reflected in different distributions of pupil size. In the current work, we examined the distribution of pupil size in humans across naturally varying levels of arousal.

Our main question was whether increasing levels of arousal were related to task-specific patterns of brain interaction in putatively different cognitive states. We observed that the level of integration and segregation changed with arousal level in some regions across tasks. In particular, regions from the salience network were more segregated at intermediate arousal and more integrated at higher arousal levels in all tasks. This is consistent with the direct connections between the saliency and novelty detection system and LC (Corbetta et al., 2008). Previously, a bilateral region showing significantly larger pupil-participation coefficient correlation after administration of atomoxetine, a blocker of NE transporter (mimicking a high arousal state), was located close to the insula, a center within the salience network (Figure 1d in Shine et al. 2018b). Activity in the salience network also correlated with pupil size during resting state (Schneider et al., 2016). Taken together, these results suggest that arousal is tightly linked to the topological properties of network integration of the salience network. In addition, we observed some specificity in the different tasks. Although overall brain patterns (**Figure 2**) did not entirely match the activation patterns associated to these tasks (Fox et al., 2005; Mäki-Marttunen et al., 2019; Mäki-Marttunen et al., 2020b), the inferior frontal gyrus in AX-CPT (i.e. a cognitive control task), and frontal-eyes fields in MOT (i.e. a dynamic visual attention task) significantly increased integration with arousal, which may hint that these effects may be in part related to task-related activity. Taken together, our results support the view that pupil size is a marker of ongoing network states (Schwalm & Jubal, 2017), and point towards arousal as one mechanism affecting brain coordination between specific areas in tasks involving different cognitive states.

When looking at the average trend of integration and segregation in the different tasks, we found a U-shaped pattern with arousal (**Figure 3**). Arousal led to increasing integration at larger arousal levels, but also specific effects become evident at the lowest arousal level. A non-linear pattern between neuromodulatory function and brain or behavior is a common finding (Aston-Jones & Cohen, 2005; Cools & D’Esposito, 2011; Mittner et al., 2016; Shine, 2019; Shine et al., 2018b). Several previous studies have assessed the relation between brain integration and pupil size or levels of catecholamines in humans. Overall, the studies showed an increased brain integration with increased arousal (Pfeffer et al., 2020; Shine et al., 2016; van den Brink et al., 2018), in agreement with animal studies where chemogenetic activation of LC led to larger brain integration (Zerbi et al., 2019). Some studies, on the other hand, presented opposite findings, with increased brain segregation at elevated levels of cathecolamines (Guedj et al., 2017b; van den Brink et al., 2016b), or no change (Pfeffer et al., 2020) during resting state. These contradicting findings could be due to the fact that pharmacological manipulations affect several neuromodulatory systems at a time, and that they may affect neuromodulatory receptors with opposing effects in brain circuitry. On the other hand, our results of a non-linear relation between arousal and brain integration, suggest that the effects of arousal on brain coordination may be more complex than a linear relation. Investigating more extreme arousal states than the ones covered here (e.g. stress or drowsiness) with a similar approach than the one we used may help describe the relation between arousal and brain effects more extensively.

How to explain the observed relation between arousal and brain topology? Animal studies show that increases in arousal (as indicated by larger pupils) are associated to suppression of slow oscillations, cortical desynchronization and cortical activation (Gervasoni et al., 2004; McGinley et al., 2015b; Takahashi et al., 2015). This accompanies active behavior such as locomotion and whisking, thus being required for accurate processing of incoming stimuli (for a review see Schwalm and Jubal 2017). A recent paper in humans showed that the wake state is characterized by larger brain integration as compared to states of low arousal (such as sleep), and relates to specific patterns of global and local synchronization, as well as the degree of correlation between functional to structural connectivity (Hahn et al., 2020). Further studies relating these electrophysiological signatures to fluctuations in arousal in the wake state will allow relating circuit to network properties of the brain as a function of arousal.

We observed specific regions showing a significant pattern of integration with arousal in each task. One possibility is that different “pupil-indexed” network states (Schwalm & Jubal, 2017) may be enabling different brain circuits during the different tasks. With their wide projections, neuromodulatory systems increase neural gain at the brain level, but this generalized effect interacts with local processes engaged in the specific task manipulations (Mather et al., 2016; Shine et al., 2018b; Stitt et al., 2018). Such local processes would be local synaptic excitatory and inhibitory processes and inter-regional coupling. In addition, we have not distinguished between on-task periods and off-task periods where the mind wanders, which may have certain brain signatures (e.g. more cohesion) in specific networks but different behavioral signatures (Mittner et al., 2016). Further research with different tasks engaging different levels of arousal could further assess commonalities and differences on brain effects (e.g. learning vs. task solving, exploration vs. exploitation, focused attention vs. mind wandering, etc.). In sum, our findings are consistent with theories of arousal-related neuromodulatory systems that propose their engagement in modulating neural processes in an adaptive manner (Aston-Jones & Cohen, 2005; Zénon, 2019).

The analysis of behavior as a function of arousal level pointed to an inverted-U pattern. Even though our tasks involved different cognitive processes, and the level of accuracy differed between them, this was significant for accuracy measures in both of the tasks examined. This pattern is in close agreement with the proposal that performance relates to arousal level through the Yerkes-Dodson curve, with optimal performance at intermediate levels or arousal, and decaying performance to extremely low or extremely high levels (Aston-Jones & Cohen, 2005). Previous studies reported such a relation between pupil and performance during attentional tasks (Murphy et al., 2011; van den Brink et al., 2016a). Together with the brain results, we provide new evidence in humans that arousal level, brain coordination and behavior may be linked in a specific way for the expression of different behavioral states.

Some limitations to our study should be acknowledged. Because of the way we defined pupil size intervals, there were unequal amount of samples per category, as well as different percentage of categories per task. However, overall we were able to obtain sufficient samples for all bins in each task. In addition, the design of the tasks differed; the MOT task had longer trials than the AXCPT, while the RS was unconstrained. These differences may have affected the sustained pupil, and comparison of different tasks with similar design parameters may disentangle the effect that this has in pupil size distribution. However, we expected that the percentage of task-related pupil change is relatively small compared to the sustained effects compared here. Another caveat is that the relation between arousal level and brain measures found here does not imply directionality or causation. However, animal studies and pharmacological studies indicate that arousal is a driving force that affects the degree of interaction between neuronal circuits (McGinley et al., 2015b).

In sum, we presented evidence in humans that brain interaction follows an inverted-U pattern as a function of fluctuating arousal during different cognitive states. The salience network consistently showed this pattern, indicating a generalized effect of arousal on this network. The different cognitive tasks also showed specific regions showing a significant effect of arousal in its level of integration with the rest of the brain. We also found higher accuracy at intermediate levels of arousal in the two tasks. Our results support the role of arousal-related neuromodulatory systems coordinating the communication across brain networks in an adaptive manner, and offers opportunities to further study this effect in the healthy brain as well as in cases where neuromodulatory systems are dysfunctional.

## Ackonwledgements

I thank Prof. Thomas Espeseth for facilitating the dataset used for this study. I thank Dr. Thomas Hagen and Michelle Antal for their contribution in the data acquisition of the data.

